# Foraging decisions of snowshoe hares in response to experimentally induced coat-colour mismatch

**DOI:** 10.1101/2023.05.05.539566

**Authors:** Joanie L. Kennah, Michael J. L. Peers, Juliana Balluffi-Fry, Isabella C. Richmond, Eric Vander Wal

## Abstract

Animals may exhibit various strategies to mitigate the adverse effects of phenological mismatch. In species experiencing coat colour mismatch, the effects of lost camouflage on the susceptibility to predation may be compensated for with other antipredator traits, such as altered foraging decisions, and may further depend on the intensity of risk. We artificially simulated coat colour mismatch and predation risk in wild-caught snowshoe hares and measured their forage intake rate of black spruce browse, intraspecific selection for forage quality, i.e., % nitrogen of browse, and resulting body mass loss across different risk levels, simulated by cover or lack thereof. We found that hares did not adjust their intake rate in response to mismatch, but hares in our high-risk treatment ate significantly more than hares in our low-risk treatment. Mismatched brown hares, however, selected for more nitrogen-rich forage than their matched brown counterparts. Mismatched white hares lost 4.55% more body mass than their matched white counterparts, despite not reducing their intake rate. Hares in our high-risk treatment lost 1.29% more body mass than those in covered enclosures. We suggest that the increased selection for nitrogen-rich forage observed in brown mismatched hares may occur to mitigate the body mass loss consequences of mismatch. Similarly, the increased intake rate of hares in clear roof enclosures relative to those in opaque roof enclosures may be a compensatory behavioural response to increased body mass loss. Our results highlight the potential indirect effects of coat colour mismatch on snowshoe hares, but also the corresponding behavioural mechanisms that may partially mitigate these effects.

## INTRODUCTION

Many prey traits have evolved to mitigate the risk of predation within and across ecological contexts (Endler 1995). For example, background matching is an antipredator trait where animals resemble the general colour or pattern of their environment (Caro 2005, Skelhorn and Rowe 2015, Duarte et al. 2017). In seasonal environments, some species achieve this camouflage through colour polymorphism, where they change colour biannually in the autumn and spring to match the presence or absence of snow (Zimova et al. 2018, Bernardi et al. 2021). The plumage or coat moults that allow this seasonal colour change are often driven by changes in photoperiod (Zimova et al. 2014, Funakoshi et al. 2017). Climate change is causing increasingly early springs and decreased snow duration (Brown and Mote 2009), which results in mistimed coat and plumage colour change with snow cover in many bird and mammal species, e.g., mountain hare (*Lepus timidus*), snowshoe hare (*Lepus americanus*) and Alpine rock ptarmigan (*Lagopus muta helvetica*) (Imperio et al. 2013, Pedersen et al. 2017, Zimova et al. 2018). Mismatched coat or plumage colour is presumed to increase mortality risk in many species (Imperio et al. 2013, Pedersen et al. 2017, Kumar et al. 2020). However, prey species exhibit suites of antipredator traits that act in concert to reduce predation risk (Makin et al. 2017, Kennah et al. 2023). Whether mismatched animals could potentially adjust other antipredator traits to reduce any negative survival costs driven by lost camouflage remains unknown.

Vigilance and foraging decisions are antipredator behaviours that animals can adjust to balance the trade-off between food acquisition and predation risk (Winnie and Creel 2017). Prey live in heterogeneous landscapes of risk and resources, and their decisions to allocate time to foraging or vigilance behaviours intrinsically affect their nutritional intake, energetic expenditure and susceptibility to predation (Hodges and Sinclair 2005, Gaynor et al. 2019, Richmond et al. 2022; Shiratsuru et al. 2023). Different foraging outcomes, such as changes in total intake rate and selection of forage, may result from increased vigilance and may be used to achieve optimal energetic gains under different risk scenarios (Verdolin 2006). Animals may adjust their intake rate by modifying decisions such as how long to consume forage, how long to search for forage, and how long to handle forage. These fine-scale foraging decisions that animals make to balance energy gain under different risk scenarios are predicted under the Optimal Foraging Theory (OFT) (Charnov 1976, Pyke et al. 1977). Similarly, according to the OFT, animals are expected to choose an optimal diet to maximize gains of a limiting nutrient or currency (e.g., energy or nitrogen) under various risk scenarios. For example, common voles (*Microtus arvalis*) decrease total food consumption with decreased feeding duration when exposed to high perceived predation risk (Eccard et al. 2020), while elk (*Cervus elaphus*) move from habitats with high-quality forage when predators are present on the landscape (Hernández and Laundré 2005). Prey may become less selective and thus experience reduced nutritional intake in high predation risk settings (Hawlena and Pérez-Mellado 2009, Hawlena and Schmitz 2010). Such foraging strategies are influenced by perceived predation risk, which varies across time and space, leading to dynamic responses by prey species altering the trade-off between food acquisition and predation risk (Brown and Kotler 2004).

Animals that perceive heightened predation risk when their camouflage is reduced may therefore adjust foraging decisions to favor protection from predation. For instance, when camouflage mismatch was experimentally simulated in jumping spiders (*Marpissa muscosa*), individuals reduced foraging intensity (Steinhoff et al. 2020). Similarly, foxes reduce their intake rate on brighter moonlit nights, when they may be more obvious to predators (Mukherjee et al. 2009). However, such altered foraging strategies represent antipredator responses that may carry downstream costs, such as body mass loss and the resulting impact on reproduction (Creel 2018). By examining how colour mismatch (i.e., lack of camouflage), influences foraging decisions under varying levels of predation risk, we can gain a more comprehensive understanding of how animals navigate complex landscapes of risk. If foraging time is reduced and replaced by increased vigilance to compensate for camouflage loss, animals may mitigate the negative costs of colour mismatch driven by climate change, although downstream effects of behavioural adjustments such as body mass loss may occur.

The snowshoe hare (*Lepus americanus*) is a keystone species of the boreal forest that has large impacts on the vertebrate community (Boutin et al. 1995; Peers et al. 2020a). Snowshoe hares change their coat colour in the autumn and spring to match the presence and absence of snow (Grange 1932). As climate change decreases the duration of the snow seasons (Danco et al. 2016), hares are becoming increasingly mismatched with their environment (Mills et al. 2013). Predation is a strong selective force in the evolution of snowshoe hares (>90% mortality attributed to predation; Peers et al. 2020b), and coat colour mismatch has been linked to decreased survival in this species (Zimova et al. 2016). However, hares can respond to high predation risk by adjusting their foraging and movement behaviour (Hutchen and Hodges 2019, Shiratsuru et al. 2021). The trade-off between food acquisition and vigilance to balance predation risk is critical for snowshoe hares as they are income breeders; that is, they do not rely on stored reserves and can die after losing 18% of their body weight (Pease et al. 1979, Kosterman et al. 2018). Therefore, the snowshoe hare is an excellent species to investigate whether foraging decisions are altered to compensate for any negative coat colour mismatch effects.

We sought to answer how coat colour mismatch affects snowshoe hare foraging decisions under different simulated perceived risk levels. We experimentally manipulated coat colour mismatch in wild snowshoe hares at different autumn coat colour change progression levels by placing them in enclosures that had simulated brown or white ground. Hares foraged during cafeteria experiments in enclosures that further provided overhead cover, or lack thereof, as a proxy of avian predation risk. We chose this proxy since hares are known to behaviourally adjust to the perceived risk of open canopy and because the majority of known hare mortalities in our system were assessed as avian (Hutchen and Hodges 2019, Richmond et al. 2022). We examined how intake rate, intraspecific selection for nutritious forage, i.e., high Nitrogen composition (%), and weight loss varied between matched and mismatched hares, and how perceived predation risk affected this difference. We hypothesized that snowshoe hares would decrease forage consumption and selection for nitrogen-rich forage when mismatched and exposed to perceived avian predation risk to increase predator avoidance at the expense of nutritional intake. As such, we predicted that mismatched hares in experimental enclosures lacking cover to simulate perceived avian predation (riskiest treatment) would have (i) the lowest total intake rate and (ii) the lowest intraspecific selection for nitrogen-rich forage. Consequently, we predicted that these modified foraging decisions would result in mismatched hares exposed to perceived avian predation risk experiencing (iii) the most body mass loss during experimental trials. Understanding whether animals may – or may not – respond to colour mismatch is vital to predict their ability to persist in the face of increased environmental stochasticity.

## MATERIAL & METHODS

### Study area and snowshoe hare trapping

Our study occurred from October to December of 2020, in the central-eastern region of Newfoundland, Canada (Lat: 48.35 N, Long: −53.97 W; see Figure S1). This region is a mature white spruce plantation (*Picea glauca*) interspersed with black spruce (*Picea mariana*) and paper birch (*Betula papyrifera*). The understory is characterized by lowbush blueberry (*Vaccinium angustifolium*), sheep laurel (*Kalmia angustifolia*), and Labrador tea (*Rhododendron groenlandicum*). Snowshoe hares at this study site experience avian predation by great horned owls (*Bubo virginianus*) and goshawks (*Accipiter gentilis*), as well as terrestrial predation by Canada lynx (*Lynx canadensis*) and coyotes (*Canis latrans*).

We live trapped snowshoe hares on a 500 m by 500 m trapping grid. The trapping grid contains 50 live traps (Tomahawk Live Trap Co. Tomahawk, WI, USA) that are separated ∼ 37-75 m along six transects. We set traps at sunset and baited them with fresh apple slices, rabbit chow, and alfalfa cubes. We checked traps and handled hares within 12 hours of baiting and setting, and installed eartags on captured individuals to identify them upon subsequent recapture. We recorded the mass (to the nearest 0.02 kg), sex, right hind foot length (cm), and age class of each individual captured, and identified 48 unique individuals during our study.

### Experimental setup

We held hares captive in experimental enclosures that allowed us to evaluate their foraging decisions under different coat colour mismatch and perceived predation risk treatments. Our eight enclosures were 100 cm wide, 90 cm high, and 120 cm long. We installed enclosures at least 10 m apart in areas lacking overhead cover, but with horizontal forest cover to obscure neighbouring enclosures. We created an opportunity for cover access with a hutch secured in each enclosure (Figure S2) and installed camera traps (Reconyx Hyperfire) facing the enclosures to collect temperature (°C) and photo data every 5 minutes during trials. We simulated perceived avian predation risk by covering the roofs of four enclosures with transparent plexiglass sheets (Figure S2a-d), thus allowing overhead visibility while protecting hares from precipitation. Conversely, we covered the roofs of four control enclosures with opaque tarps to simulate perceived protection from avian predation risk (Figure S2e-h). Indeed, prey have been shown to display stronger behavioural adjustments to habitat characteristics, such as vegetative cover or lack thereof, than cues of live predators (Verdolin 2006). More specifically, snowshoe hares are known to behaviourally adjust to the perceived risk of open canopy (Hutchen and Hodges 2019). We placed three iButtons (iButton Thermochron) in one opaque roof enclosure and another three iButtons in a transparent roof enclosure to verify that roof type did not influence enclosure temperature (Figure S3). We experimentally simulated coat colour mismatch by placing hares at different levels of coat colour change on white or brown enclosure floors (Figure S2).

We selected hares weighing > 1300 g for experimental trials. We repeated experimental trials for hares if they had recovered to within 5% of their original mass and a minimum of 10 days had elapsed since their previous trial. We did not allow hares to complete more than four experimental trials during our field season. Hares were held captive for a maximum of 72 hours, during which they underwent an adjustment period, two feeding trials and a recovery period (detailed below). Snowshoe hare handling and experimental trials were approved by Memorial University of Newfoundland’s Animal Care and Use Committee (Protocol #18-02-EV).

### Feeding trials

We evaluated how our different treatments influenced total browse consumed by snowshoe hares, but also intraspecific selection for browse quality, since animals are known to adjust their selection for forage quality with perceived predation risk (Edwards 1983, Hernández and Laundré 2005, Barnier et al. 2014). We chose to test selection for quality using naturally occurring quality variation within a single browse species. Indeed, intraspecific forage decisions are likely prominent for herbivores in low plant diversity systems, such as the boreal forest (Balluffi-Fry et al. 2022). Black spruce was used to test intraspecific foraging decisions because it is widespread across our study site and constitutes the majority of hare diet in Newfoundland after the senescence of deciduous plants, thus it represents a realistic diet item considering the timing of our study (Dodds 1960, Rodgers and Sinclair 1997). We chose % nitrogen (N) content as our proxy for black spruce quality, as it is known as a limiting item for herbivores (Fagan et al. 2002, Boersma et al. 2008). Further, selection for black spruce with high N content has been shown to mitigate snowshoe hare weight loss in our system (Balluffi-Fry et al. 2021; See supplement for detailed spruce sampling method).

We offered 100 g piles of both high-quality and low-quality black spruce boughs to snowshoe hares during 15-hour feeding trials that occurred between 18:00 and 9:00, to reflect the hours during which hares are most active (Studd et al. 2019; See supplement for detailed spruce samping methods). We presented each pile in a basket on either side of the enclosure (Figure S2), and we randomly selected which side to place high and low quality forage. We measured total browse consumption by collecting the remaining spruce after feeding trials and subtracting its mass from the total spruce offered, i.e., 200g. Finally, we assessed how feeding decisions affected snowshoe hare body condition by measuring mass change during these 15-hour feeding trials. Each hare underwent one to two feeding trials in the context of our 72-hour experimental trials, which consisted of four distinct trial phases (Figure S4).

### 72 – hour experimental trail phases

To begin, each hare was first randomly assigned to an enclosure with a clear or opaque roof, and a brown or white floor. The first experimental trial phase was a 9-hour adjustment period (9:00 to 18:00, Day 1) during which we provided hares with water *ad libitum*, an apple slice, rabbit chow ad libitum and ∼ 200 g of mixed quality black spruce. We created the adjustment period to ensure that all hares began their feeding trial after getting the opportunity to feed to satiation and to allow hares to habituate to enclosures. The second phase (18:00 Day 1 to 9:00 Day 2) was a 15-hour feeding trial, where hares received separated piles of 100 g of high quality and 100 g of low-quality spruce and water ad libitum. For the third phase (9:00 Day 2 to 18:00 Day 3), the recovery period, hares were transferred to a randomly selected new enclosure that provided them with the opposite perceived predation risk treatment, i.e., clear or opaque enclosure roof, as their first food choice trial but the same mismatch treatment, i.e., floor colour. This recovery period is a necessary phase within the 72-hour experimental trials, as hares are known to lose weight during single species feeding trials that could impact foraging decisions during the next feeding trial, and survival post release (Pease et al. 1979, Rodgers and Sinclair 1997). During this recovery period, we provided hares with rabbit chow and water ad libitum. The fourth and last phase of the experimental trial (18:00 Day 3 to 9:00, Day 4) was a second 15-hour feeding trial, identical to the first, but in the new enclosures which provided the opposite perceived predation risk treatment.

To monitor mass change and ensure that hares remained within safe mass loss thresholds, we weighed them after each phase of the experimental trial (Figure S4). Hares that did not meet our requirements for safe mass loss during experimental phases were not held for the subsequent steps of the trial and were immediately released at their capture site. Mass loss thresholds were defined as: minimum mass maintenance during the adjustment phase, maximum mass loss of 8% during feeding trial phase, and mass recovered to at least within 5% of initial mass at trapping during recovery phase (Figure S4). Prior to release, we assessed coat colour of each hare by visually ranking its white coat to the nearest 10%. Coat colour rankings were all completed by the same observer, and intra-observer correlation in coat-colour rankings from photos post-hoc was high (> 0.9; Table S1). Animal handling and experimental trial procedures were approved by Memorial University’s animal use ethics committee (AUP 20-02-EV).

### Statistical analyses

We generated three linear mixed-effects models to evaluate how different mismatch and perceived predation risk treatments affected foraging decisions and mass change. To test prediction (i), we calculated total intake rate (g spruce/kg hare) as the difference between the sum of all spruce offered and remaining at the end of the 15-hour feeding trials (g), divided by individual hare mass prior to commencing feeding trials (kg). To test prediction (ii), we calculated intraspecific selection for quality as the difference between high-quality spruce intake rate (g spruce/kg hare) and low-quality spruce intake rate (g spruce/kg hare). As such, in our intraspecific selection for quality model, positive effect sizes are associated with selection for high-quality spruce, and negative effect sizes are associated with selection for low-quality spruce. To test our final prediction (iii), weight change (%), was calculated as the percentage mass difference between each snowshoe hare’s weigh-in immediately before and after the feeding trials.

All linear mixed-effects models included the same fixed explanatory variables: average temperature during the feeding trial, mismatch, perceived predation risk, habituation, and individual ID as a random factor. We calculated the average temperature (°C) during feeding trials from a minimum of 13 hourly temperatures recorded by camera traps at each enclosure during the 15-hour feeding trials. We chose to include temperature in our models, as snowshoe hares are known to increase their intake rate as it gets colder (Sinclair et al. 1982, Balluffi-Fry et al. 2021), which may subsequently affect weight change and intraspecific selection for quality. Coat colour mismatch was defined as the difference between enclosure floor % white, i.e., 0% for brown floors or 100% for white floors, and snowshoe hare coat % white. We first included mismatch in our models as a binary variable, i.e., mismatched or not, disregarding whether a hare was mismatched white or brown. Hares displaying an absolute contrast of 60% or more with their enclosure floor were considered mismatched, as per previous studies (Mills et al. 2013, Zimova et al. 2014) (Table S2). To increase the specificity of the analysis, we then modelled the effect of different mismatch categories on foraging decisions and mass change. Thus, our mismatch categories were: white mismatch, i.e., white hare on brown ground; brown mismatch, i.e., brown hare on white ground; white match, i.e., white hare on white ground; and brown match, i.e., brown hare on brown ground. We included simulated predation risk (i.e., roof type) as a binary variable. Habituation was included in our models, as hares are known to increase forage consumption when they have previously been held captive for another feeding trial within a same field season (Balluffi-Fry et al. 2021). As such, we considered habituation as a fixed binary factor for whether or not hares had undergone at least one feeding trial within our field season. We also included a random effect for individual ID to control for non-independence of data within individuals. We assessed model fit using marginal and conditional R-squared using the “r.squaredGLMM” function in the package MuMin (Nakagawa et al. 2017, Barton 2020). We carried out statistical analyses in R version 4.0.4 (2021) (R Core Team, 2021). We considered P ≤ 0.05 as our significance threshold and reported all means with ± 1 standard error.

## RESULTS

### Overview

We held 31 individuals captive for feeding trials, of which 23 met our criteria to complete full 72-hour experimental trials, whereas 8 individuals only completed a single feeding trial as they exceeded mass loss criteria to be considered for a second trial. There were 12 individuals that were recaptured throughout the field season and repeated one or more partial or full experimental trials. As a result, we completed 79 15-hr feeding trials and 32 full 72-hour experimental trials. The average daily temperature and precipitation during our study was 4.4 ± 5.1°C and 3.7 ± 6.1 mm, respectively. The hares we captured for trials ranged from 0% to 90% white (median = 20% white).

### Black spruce intake rate (prediction i)

Hares consumed an average of 96.92 ± 37.71 g of black spruce during feeding trials, and total consumption increased at colder temperatures; on average hares consumed 1.03 g/kg more for each 1 °C decrease in temperature (P=0.018; Figure 1). Contrary to our prediction, hares exposed to perceived predation risk, i.e., in clear roof enclosures, consumed significantly more spruce than those in opaque roof enclosures (P=0.023; Figure 1). For instance, when the temperature was held at its mean (4.14°C), matched hares in clear roof enclosures consumed 8.51 g/kg more than matched hares in opaque roof enclosures. There was no evidence that mismatch influenced black spruce intake rate, regardless of mismatch type (Table 1). Individuals that had already completed a feeding trial during our field season ate on average 13.56 g/kg more than those being held captive for the first time (P=0.008; Figure 1).

**Figure 1.**
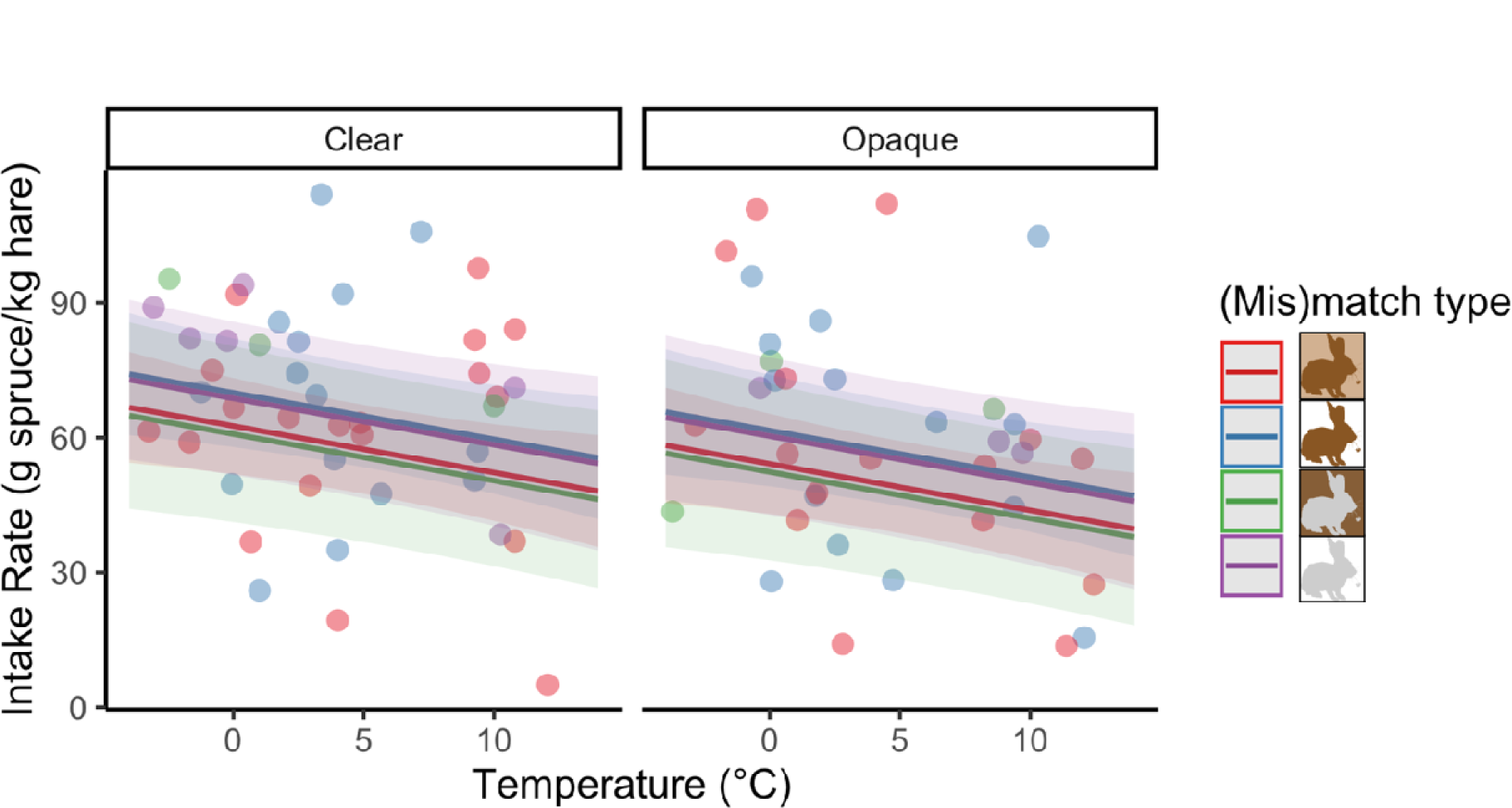
Effect of temperature on black spruce intake rate for experimentally matched or mismatched snowshoe hares under perceived predation risk treatments, i.e., clear or opaque enclosure roof. Effect is modelled for individuals during their first time held captive. Shaded areas represent 95% confidence intervals.

**Table 1.**
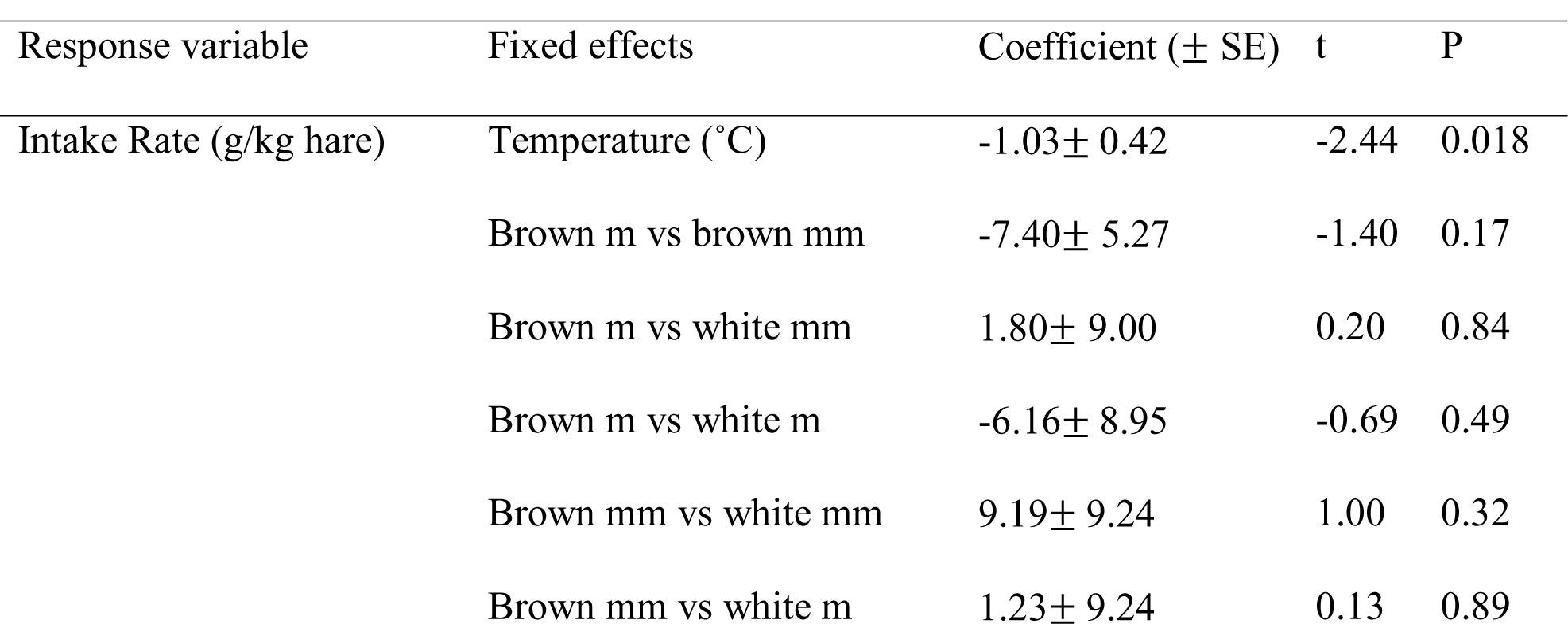
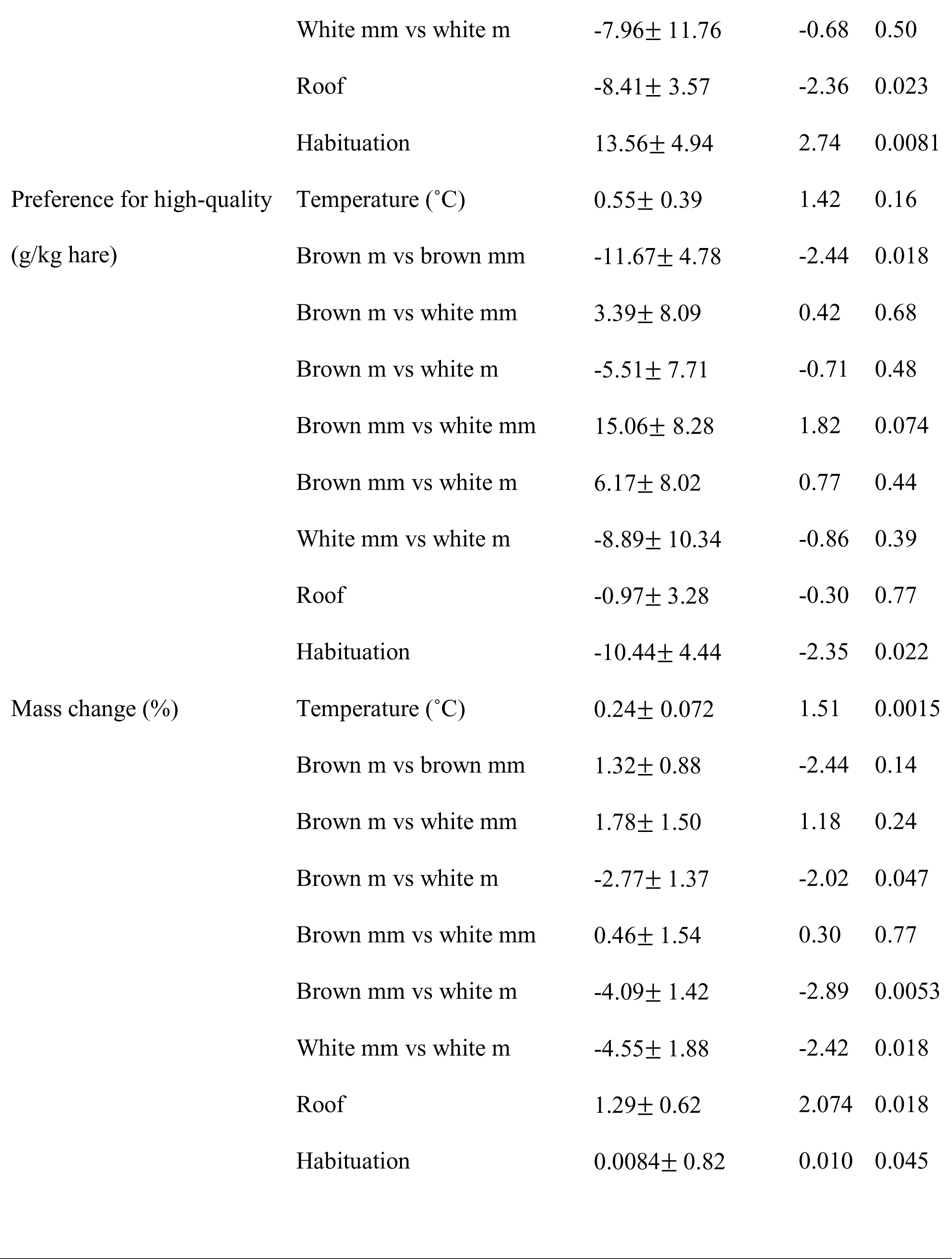
Effects of variables included in linear mixed-effects models for black spruce intake rate (marginal R^2^= 0.15, conditional R^2^=0.72), selection of high-Nitrogen spruce (marginal R^2^= 0.13, conditional R^2^=0.63), and body mass change (marginal R^2^= 0.21, conditional R^2^=0.57). In the preference for high-Nitrogen model, positive effect sizes are associated with selection for high-Nitrogen spruce, and negative effect sizes are associated with selection for low-Nitrogen spruce. In the body mass change model, positive effect sizes are associated with mass gain, whereas negative effect sizes are associated with mass loss. All models also include individual ID as a random effect. The reference variable for perceived predation risk, “roof”, is the clear roof and the reference for habituation is the first time hares were held captive. For mismatch variable: m = match; mm= mismatch.

### Intraspecific selection for nitrogen (prediction ii)

During feeding trials, hares consumed on average 50.57 ± 24.73 g of N-rich black spruce versus 47.07 ± 22.88 g of N-poor black spruce. Intraspecific selection for N-rich spruce was significantly (P = 0.02) lower during subsequent recaptures, where hares that had previously undergone experiments ate 10.44 g/kg less N-rich spruce than nitrogen-poor spruce (Figure 2). Mismatch only significantly affected black spruce nitrogen selection for brown hares, in that mismatched brown hares ate on average 11.67 g/kg more N-rich spruce than matched brown hares (P = 0.02) (Figure 2; Table 1). Intraspecific selection for N was not affected by perceived predation risk treatment (P = 0.77) or ambient temperature (P = 0.16).

**Figure 2.**
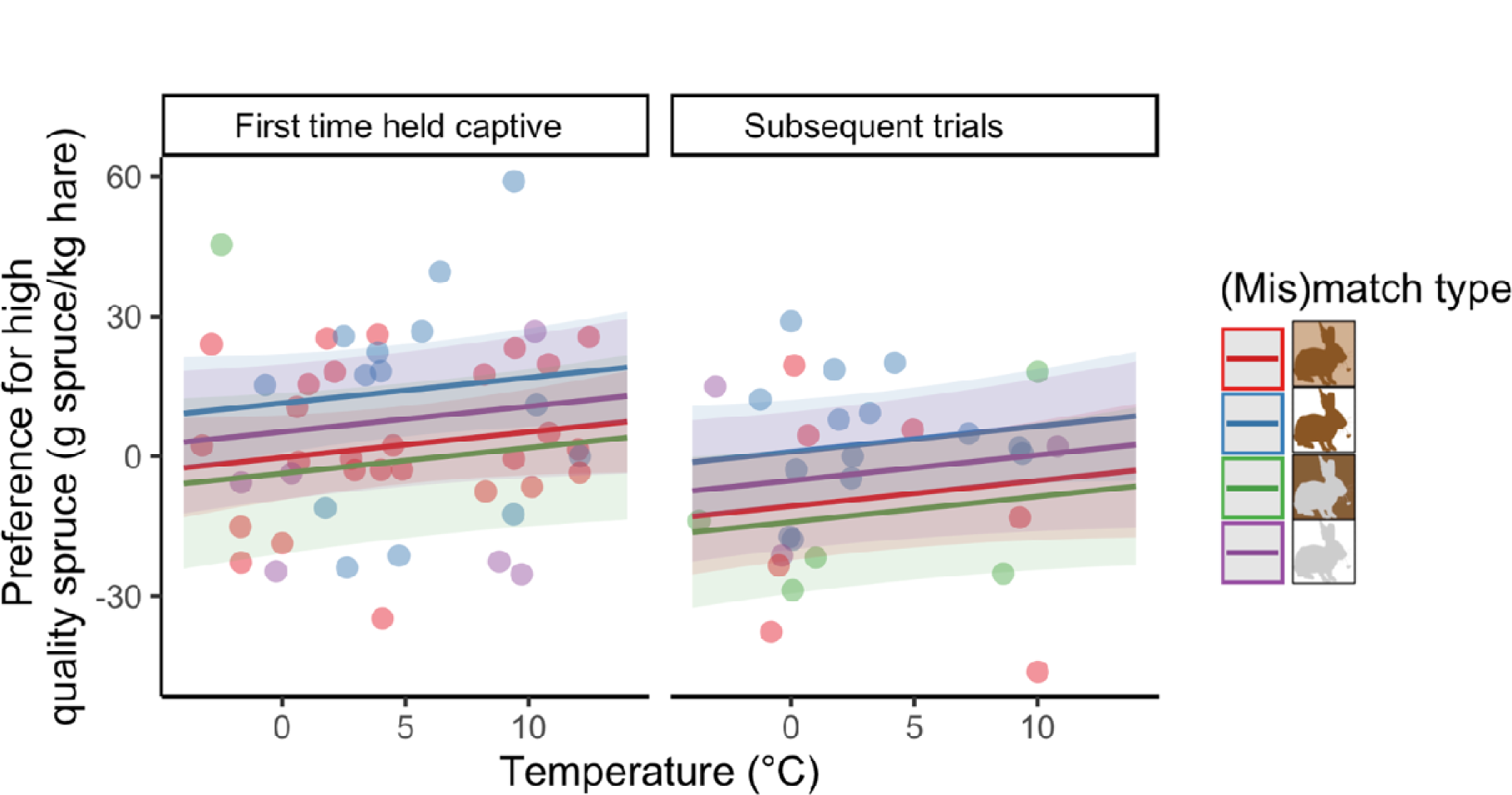
Effect of temperature on selection of N-rich black spruce, i.e., N-poor black spruce intake rate subtracted from N-rich black spruce intake rate, for experimentally matched or mismatched snowshoe hares. Effect is modelled for individuals in clear roof enclosures only. Shaded areas represent 95% confidence intervals.

### Mass change (prediction iii)

On average, hares lost weight during feeding trials (i.e., 8.03 ± 3.85% of their body mass). In contrast, individuals gained on average 4.24 ± 3.64%, and 6.34 ± 6.00% body mass during the adjustment and recovery periods, respectively. Mass loss was affected by temperature, and snowshoe hares lost on average 0.24% more body mass with every 1 °C drop in temperature (P = 0.002; Figure 3). White matched hares lost on average 4.55% less body mass than their white mismatched counterparts (P = 0.018; Figure 3). Similarly, white matched hares lost 4.09% less body mass than brown mismatched hares (P=0.005) and 2.77% less body mass than brown matched hares (P = 0.04; Figure 3). Brown mismatched hares lost on average 1.32% more body mass than their brown matched counterparts, although this effect was not significant (P = 0.14). There were no significant differences in body mass loss between brown mismatched and white mismatched hares; and between brown matched and white mismatched hares. Hares in clear roof enclosures lost on average 1.29% more body mass than those in opaque roof enclosures (P = 0.04; Figure 3).

**Figure 3.**
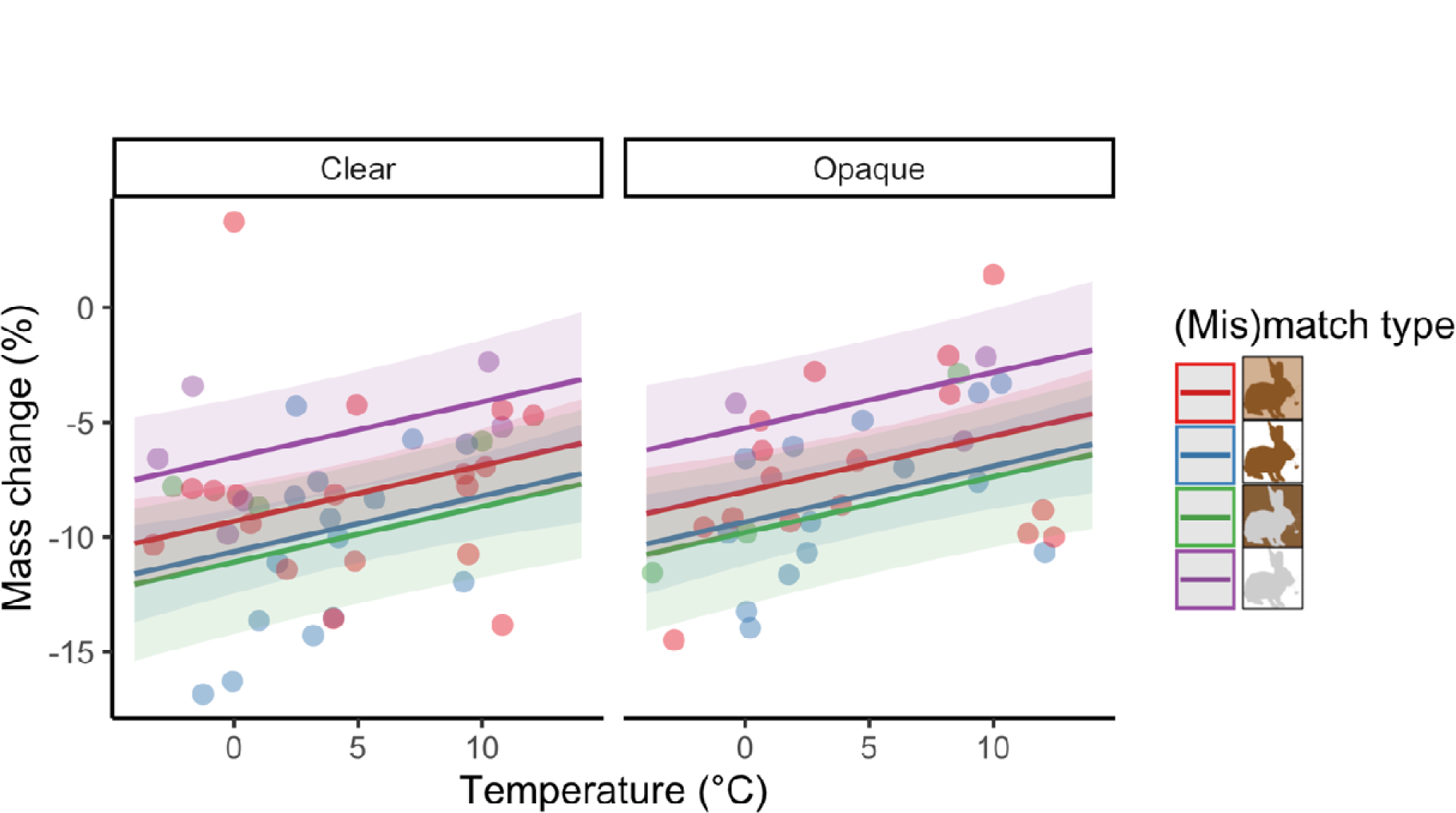
Effect of temperature on snowshoe hare mass change (%) under different mismatch and perceived predation risk, i.e., clear or opaque roof, treatments. Effect is modelled for individuals during their first time held captive. Shaded areas represent 95% confidence intervals.

## DISCUSSION

Antipredator traits function collectively to reduce an individual’s risk of being killed by predators (Garcia and Sih 2003, Kim and Velando 2015). Prey species that moult biannually to match the presence and absence of snow are expected to become increasingly colour mismatched under climate change, presumably resulting in heightened predation risk and thus increased mortality (Zimova et al. 2016, Mills et al. 2018). We sought to determine whether the lack of camouflage in snowshoe hares may be mitigated by changes in foraging decisions. Specifically, we evaluated whether coat colour mismatched snowshoe hares alter their intake rate and selection for forage N, and how these decisions correspondingly affect body mass loss. We further assessed how these decisions and resulting body mass loss were influenced by elevated risk. We found that:

1. Intake Rate: While we predicted that mismatched hares would have the lowest intake rate, we did not find any adjustments to intake rate in response to mismatch (Table 1). We predicted that hares in our elevated risk treatment, i.e., clear roof enclosures would have the lowest intake rate, but found that they unexpectedly ate more, rather than less, than hares in opaque roof enclosures (Figure 1).
2. Selection for N: We predicted that mismatched hares would have the lowest intraspecific selection for N-rich forage, but found that mismatched brown hares had a higher preference for N-rich forage than matched brown hares (Figure 2). Hares did not alter their selection for N as a response to elevated risk, which does not support our initial prediction that hares in clear roof enclosures would have the lowest intraspecific selection for N-rich forage (Table 1).
3. Body Mass Loss: As predicted, mismatched hares lost more body mass than matched hares and this effect was further amplified by heightened predation risk, as hares in clear roof enclosures lost more body mass than those in opaque roof enclosures (Figure 3; Table 1).

Although mismatched hares in our elevated risk treatment did indeed lose more body mass as expected, this effect was paired with foraging decisions we would not expect to minimize risk at the expense of nutritional intake. For instance, hares increased intake rate in response to lack of cover, and increased selection for N in response to brown mismatch, which presumably would have resulted in nutritional intake being favoured over vigilance. Thus, we suggest that mismatch generates indirect costs that are not fully compensated by modified foraging decisions.

The increased body mass loss that occurred in mismatched hares (Figure 3) may be a deleterious indirect effect of coat colour mismatch. Prey are known to respond to predation risk through the “stress response”, whereby the hypothalamic-pituitary-adrenal axis is activated and glucocorticoids are secreted to mobilize energy and face imminent threat (Boonstra et al. 1998, Sheriff et al. 2011). When the stress response is activated, physiological responses including increased body mass loss occur as a function of gluconeogenesis (Carlsen et al. 1999, Pérez-Tris et al. 2004, Hodges et al. 2006). As such, we propose that hares may associate mismatch with increased perceived predation risk, and that the resulting stress response could be a potential driver for the heightened body mass loss we observed in mismatched hares (Figure 3, Table S3). As such, coat colour mismatch may have indirect costs on individual snowshoe hares beyond the heightened mortality risk previously documented (Zimova et al. 2016, Wilson et al. 2018). Similar to mismatch, we propose that hares in our elevated risk treatment, i.e., clear roof enclosures, elicited this stress response that also exacerbated body mass loss (Figure 3), despite the increased intake rate of hares in this treatment (Figure 1). Indeed, hares experience exacerbated body mass loss with heightened predation risk (Majchrzak et al. 2022) and lack of cover has been associated with higher stress levels in some species (Kordosky et al. 2021). However, our results suggest that the increased intake rate of hares in clear roof enclosures did not compensate for potential exacerbated body mass loss due to simulated heightened predation risk.

Protein is limiting for herbivores in the boreal forest (Hodges and Sinclair 2003), and when snowshoe hares do not consume enough protein, they lose body mass (Sinclair et al. 1982, Rodgers and Sinclair 1997). Hares may mitigate body mass loss by increasing their selection for high N black spruce (Balluffi-Fry et al. 2021). We found that mismatched white hares lost 4.55% more body mass than their matched white counterparts, and that mismatched brown hares lost 1.32% more body mass than matched brown hares, although the latter was not significant (Figure 3; Table 1). Intraspecific selection for N-rich black spruce was unexpectedly greater for brown mismatched hares than for brown matched hares (Figure 2). We suggest that brown mismatched hares may increase their selection for N-rich black spruce to potentially mitigate the heightened body mass loss that occurred with coat colour mismatch. Indeed, animals adjust their diet quality selection to cope with dynamic metabolic and energetic needs (Mellado et al. 2005).

Compensatory responses to long-term predation risk have been shown to offset the adverse effects that prey experience due to the physiological responses they mount to face short-term predation risk (Thaler et al. 2012). These compensatory responses may be physiological or behavioural in nature. For example, damselfly (*Enallagma cyathigerum*) larvae reduce energy storage when exposed to predation risk (Van Dievel et al. 2016), while tobacco hornworm caterpillars (*Manduca sexta*) demonstrate compensatory feeding by increasing their intake rate after a certain level of predator exposure (Thaler et al. 2012). While we predicted that hares in our elevated risk treatment, i.e., clear roof enclosure, would decrease their intake rate in contrast with those in opaque roof enclosures, we found the opposite (Figure 1). This increased intake rate in response to lack of overhead cover contradicts previous findings (Kotler et al. 1991, Mohr et al. 2003), although these studies did not assess the impacts of overhead cover on body mass loss. We reconcile our result with our proposed interpretation of increased body mass loss as a function of increased stress. Indeed, we suggest that hares may have increased their intake rate (Figure 1) to partially compensate for the heightened body mass loss they experienced in clear roof enclosures (Figure 3). A study that manipulated predation risk to assess the physiological responses of snowshoe hares found expected physiological evidence of a stress response, but this stress response was not associated with the expected downstream consequences on body condition (Boudreau et al. 2019). Authors from this study propose compensatory behavioural or metabolic responses as potential explanations for this result. Although we did not explicitly evaluate the impacts of our experimental treatments on snowshoe hare stress response, we found increased intake rate in snowshoe hares in our high-risk treatment. As such, we suggest that hares in our study may have employed compensatory feeding through increased intake rate in an attempt to mitigate the body mass loss impacts they experienced in high-risk enclosures.

Across all mismatch and heightened risk categories, white mismatched hares in clear roof enclosures lost the most body mass (Table S3). Snowshoe hares undergo seasonal acclimatization that includes changes in coat characteristics that ultimately result in winter white hares having a lower metabolic rate than summer brown hares to facilitate winter survival (Sheriff et al. 2009a, 2009b). As a function of these changes, daily foraging time is lower for white mismatched hares than for brown hares when temperatures are below −3°C, presumably due to reduced energetic demands (Kennah et al., 2023). In this study, we found that white mismatched hares lost more weight than brown mismatched hares, implying that white coats did not confer energetic benefits (Table S3). Our results may be explained by the relatively warm temperatures experienced by mismatched hares during our feeding trials, i.e., on average 4.15°C, which were outside of the temperature range where energetic benefits were suggested in white winter-acclimatized hares (Kennah et al. 2023). This supports the idea that coat colour mismatch effects are temperature dependent, with potential indirect costs on body mass for populations in the southern parts of their range.

Coat colour mismatch has the potential to negatively impact over 21 bird and mammal species that change colour biannually, many of which are prey species that depend on camouflage to reduce their predation risk (Mills et al. 2018, Zimova et al. 2018). We tested whether coat colour mismatched snowshoe hares may be able to buffer these effects by altering their foraging decisions. We found that brown mismatched hares selected for N-rich forage, and that both brown and white mismatched hares lost more body mass than their matched counterparts. Complementary to the direct impacts of coat colour mismatch on survival that have been documented (Zimova et al. 2016, Atmeh et al. 2018, Kennah et al. 2023), our results highlight indirect mismatch effects that may additively affect fitness. We found evidence for altered foraging decisions, which only partially mitigate the consequences of increased risk and mismatch on body mass loss. We found that white mismatched hares lost the most body mass, in contrast with other mismatch categories. Our work adds to a growing body of literature that expands our understanding of how rapidly changing climate affects animals.

## ACKNOWLEDGEMENTS

We thank J. Hendrix, G. Riefesel, Y. Majchrzak, A. Robitaille, and K. d’Entremont for their help with data collection. Thank you to S. Leroux, Y. Wiersma, and the Wildlife Evolutionary Ecology laboratory and Terrestrial Ecosystem Research Group at Memorial University of Newfoundland for comments on earlier versions of this manuscript. We also thank Alec Robitaille for assistance and help with statistical analyses and R code. This work was supported by the Natural Sciences and Engineering Research Council of Canada, Northern Studies Training Program, and the W. Garfield Weston Foundation.

