## Supplementary for "Foraging decisions of snowshoe hares in response to experimentally induced coat-colour mismatch"

**This Supplementary Information includes:**

Supplementary Figures S1-S4

Supplementary Tables S1-S3

Supplementary Methods

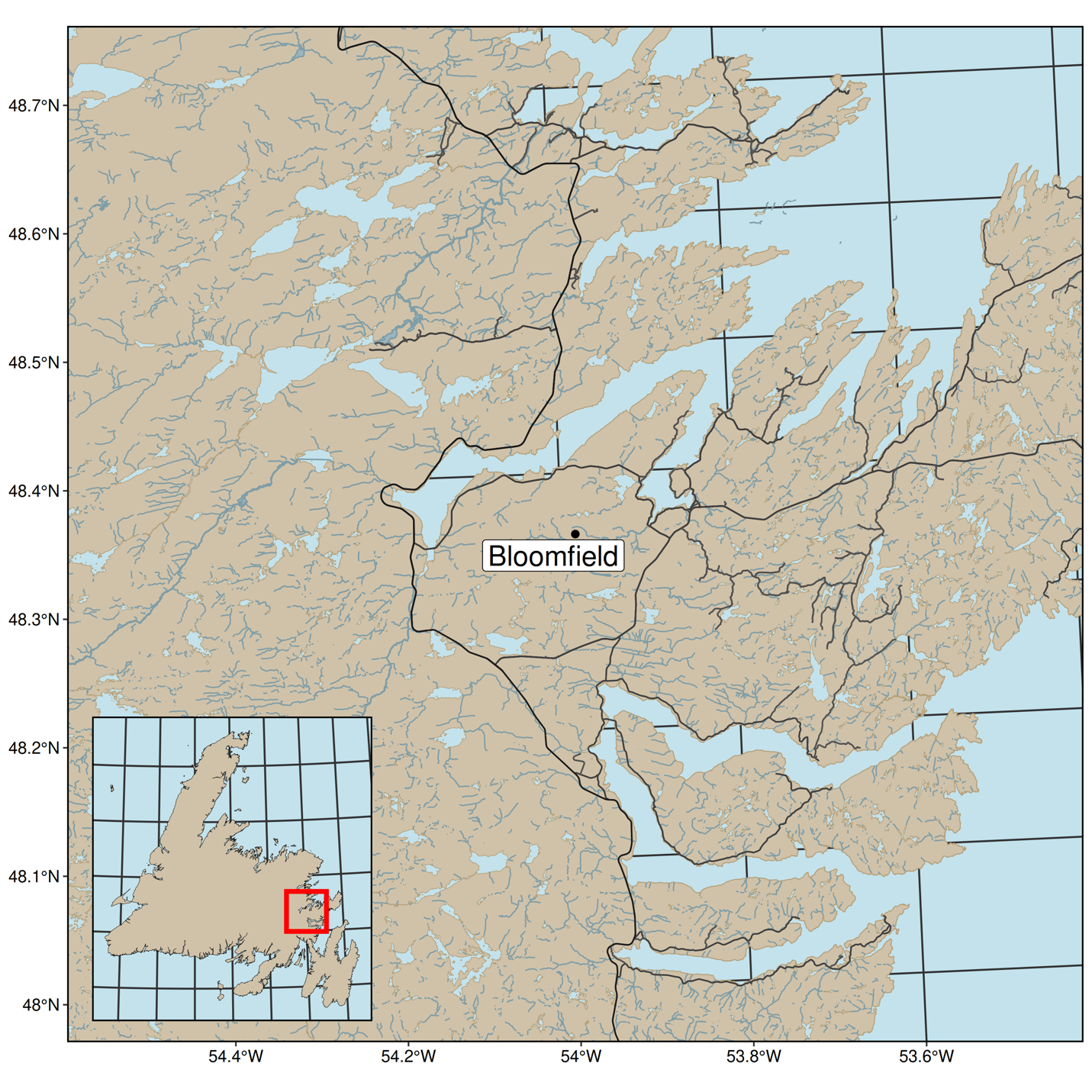

Figure S1. Our study site in Bloomfield, NL. Map is created by Alec L. Robitaille, Isabella C. Richmond and Juliana Balluffi-Fry, with roads from Open Street Map (Robitaille et al. 2021).

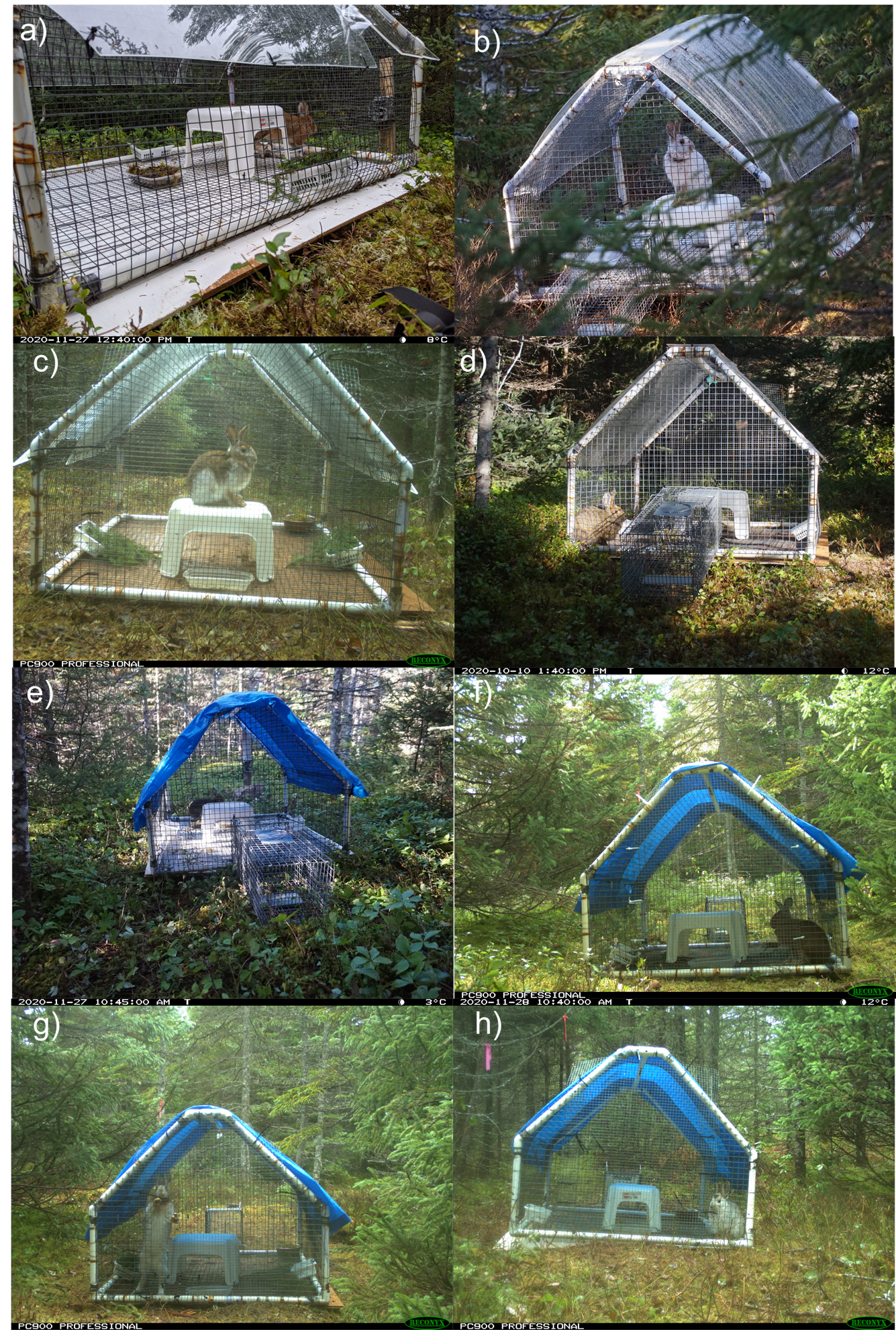

Figure S2. Experimental enclosures showing various mismatch and perceived predation risk treatments: a) mismatched brown hare with perceived predation risk; b) matched white hare with perceived predation risk; c) mismatched changing hare with perceived predation risk; d) matched brown hare with perceived predation risk; e) mismatched brown hare with perceived protection from predation risk; f) matched brown hare with perceived protection from predation risk; g) mismatched white hare with perceived protection from predation risk; h) matched white hare with perceived protection from predation risk.

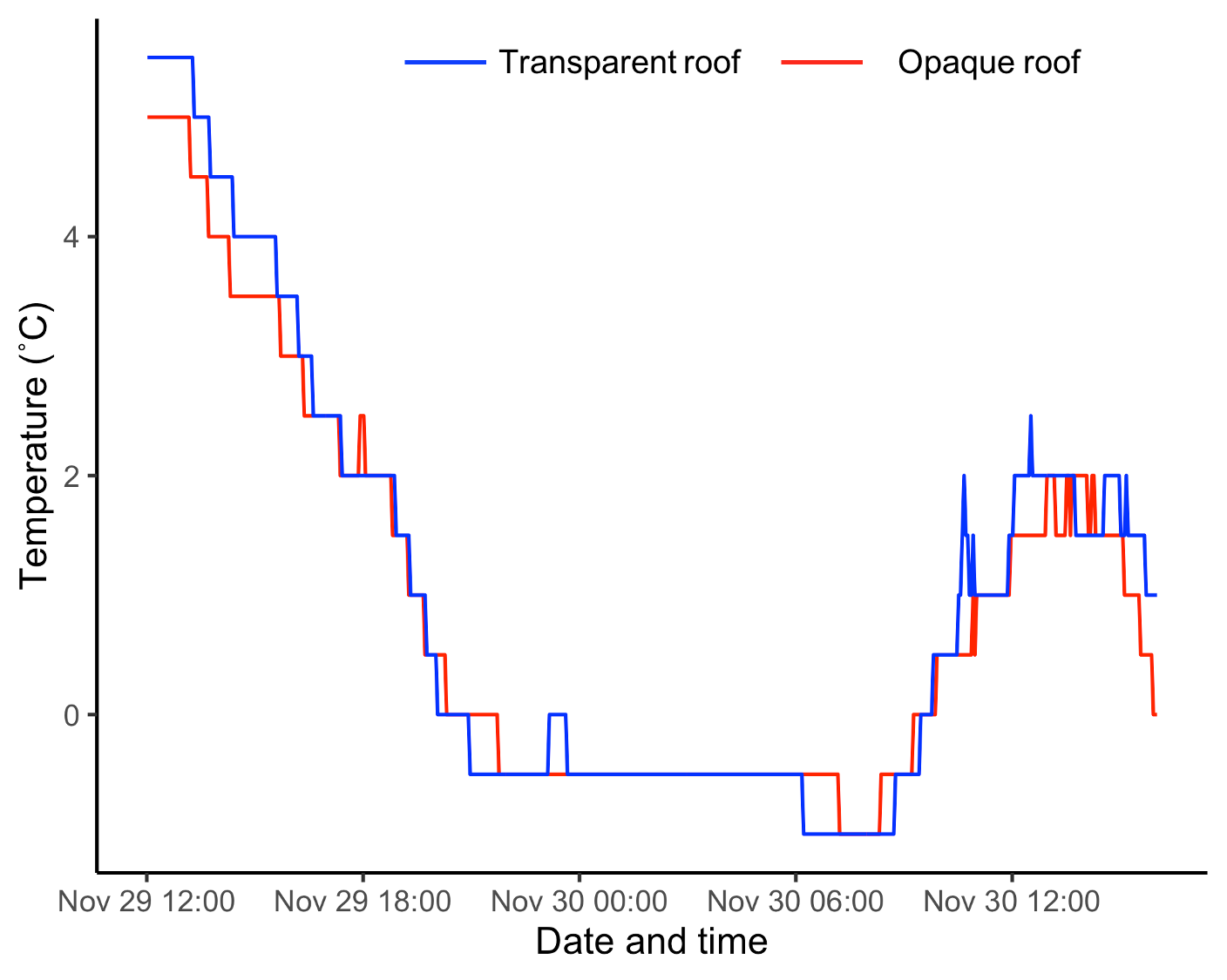

Figure S3. Temperature profile comparing clear (i.e. perceived predation risk treatment) and opaque (i.e., perceived protection from predation risk treatment) roof enclosures.

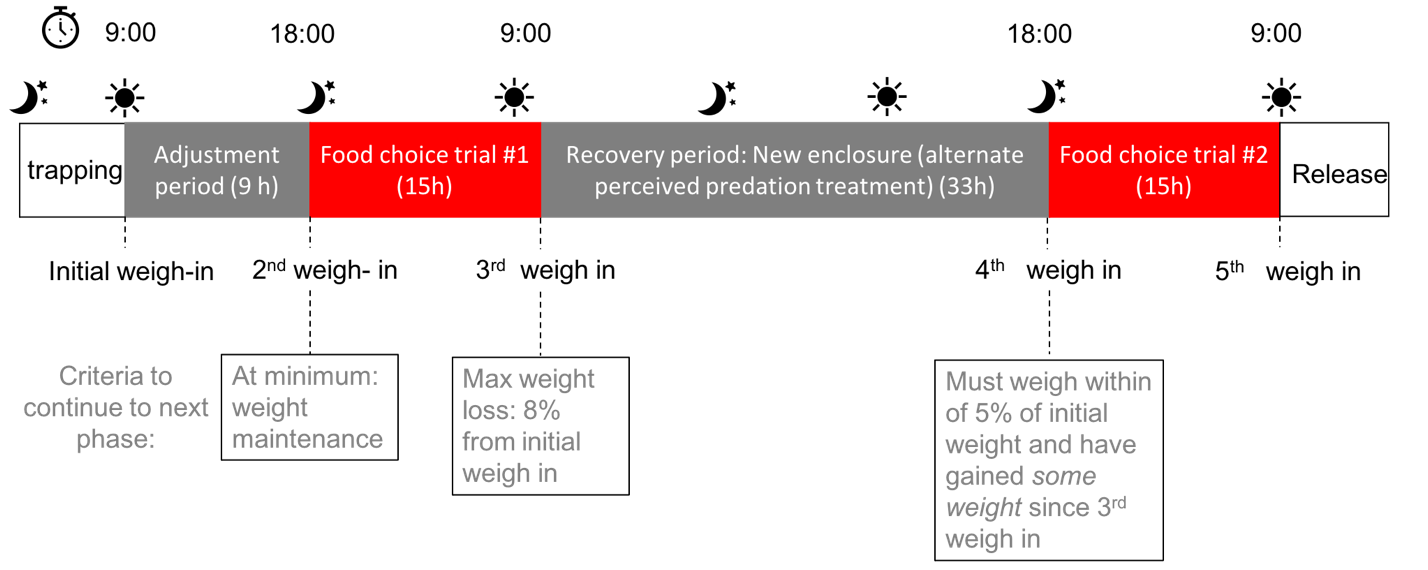

Figure S4. Outline of the phases included in our 72- hour experimental trials. Weight change threshold criteria used to determine whether hares qualified to continue onto the next phase are indicated.

Table S1. Intraclass correlation coefficients (ICC) describing intra-observer repeatability for our single consistent observer. The observer ranked coat colour twice to the nearest 5% for 31 hares (total of n=62 rankings).

|  | ICC | 95% confidence intervals | F (df1, df2) | p-value |
| --- | --- | --- | --- | --- |
| Observer 1 | 0.927 | 0.872, 0.958 | 26.2 (46,47) | < 0.001 |

Table S2. Effects of variables included in linear mixed-effects models for black spruce intake rate, selection of high-quality spruce, and body mass change when mismatch is considered as a binary variable. In the preference for high quality model, positive effect sizes are associated with selection for high-quality spruce, and negative effect sizes are associated with selection for low-quality spruce. In the body mass change model, positive effect sizes are associated with mass gain, whereas negative effect sizes are associated with mass loss. All models also include individual ID as a random effect. The reference level for predation risk, “roof”, variable is the clear roof, the reference for habituation is the first-time hares were held captive, and the reference level for mismatch is matched.

| Response variable | Fixed effects | Coefficient ($\pm$ SE) | t | P |
| --- | --- | --- | --- | --- |
| Intake Rate (g/kg hare) | Temperature (˚C) | -1.11$\pm$0.42 | -2.62 | 0.11 |
|  | Mismatch | 4.76$\pm$4.74 | 1.00 | 0.32 |
|  | Roof | -8.51$\pm$3.59 | -2.37 | 0.02 |
|  | Habituation | 12.80$\pm$4.67 | 2.74 | 0.008 |
| Preference for high-quality (g/kg hare) | Temperature (˚C) | 0.44$\pm$0.40 | 1.09 | 0.28 |
|  | Mismatch | 8.77$\pm$4.41 | 1.99 | 0.05 |
|  | Roof | -1.30$\pm$3.48 | -0.38 | 0.71 |
|  | Habituation | -12.30$\pm$4.40 | -2.80 | 0.007 |
| Mass change (%) | Temperature (˚C) | 0.23$\pm$0.07 | 3.09 | 0.003 |
|  | Mismatch | -1.87$\pm$0.79 | -2.35 | 0.02 |
|  | Roof | 1.23$\pm$0.63 | 1.95 | 0.06 |
|  | Habituation | 0.20$\pm$0.80 | 0.26 | 0.80 |

Table S3. Predicted average body mass loss (%) and 95% Confidence Intervals during a single feeding trial across simulated risk levels, i.e., match or mismatch and clear or opaque top enclosure, when temperature is held constant at its mean, and hares are in their first experimental trial (non-habituated). Letters show mismatch category pairs that are associated by a significant difference. 95% Confidence Intervals are presented in brackets for each predicted body mass loss value.

|  | Enclosure roof | | |
| --- | --- | --- | --- |
| Simulated match/mismatch type | Clear |  | Opaque |
| White match^a,b,c^ | 5.53 (3.01, 8.04) |  | 4.24 (1.63, 6.84) |
| Brown match^a^ | 8.30 (6.87, 9.73) |  | 7.01 (5.51, 8.50) |
| Brown mismatch^b^ | 9.62 (7.93, 11.32) |  | 8.33 (6.60, 10.07) |
| White mismatch^c^ | 10.08 (7.05, 13.1) |  | 8.79 (5.75, 11.83) |

**SUPPLEMENTARY METHODS**

***Spruce sampling and intraspecific quality assessment***

*Sampling location*

The black spruce used to test intraspecific selection for forage quality under our different experimental treatments was collected from a similar snowshoe hare trapping grid, “Unicorn”. The Unicorn grid is 32 km away from the grid used in our study, just north of Terra Nova National Park. This grid was chosen for spruce harvesting because its vegetative composition is similar to our grid, black spruce trees at our trapping location had already been extensively clipped for previous feeding trial experiments, and since spruce tree elemental composition data at this location was available (Heckford et al. 2021).

*Sampling to establish stoichiometric distribution maps*

Black spruce was collected from 50 sample locations at the Unicorn grid along six trapping transect lines between July and August of both 2016 and 2017 (see Heckford et al. 2021). At each sampling location, the terminal ends, i.e., new growth, of black spruce branches on juvenile trees, i.e., <2 m in height, were collected. Sampling was completed within an 11.3 m radius plot around each sampling location, in four intercardinal directions, i.e. NE, NW, SE, SW, moving clockwise until 20 g was collected in total. The new growth of black spruce branches is representative of what is typically consumed by snowshoe hares. Samples were stored at -20°C until processing. Elemental analysis of these samples was conducted at the Agriculture and Food Lab at the University of Guelph, where percent nitrogen (N) was assessed using an Elementar Vario Macro Cube. Stoichiometric distribution models were used to extrapolate N composition values across the entire Unicorn grid (Heckford et al. 2021).

*Spruce harvested for feeding trials*

We collected black spruce from seven locations that had previously been sampled for elemental composition. We determined sampling locations from the N “hot spots” and “cold spots” from previously established stoichiometric distribution maps. N content (%) at high-quality sampling locations was on average 0.93 and N content (%) at low-quality sampling locations was on average 0.87. We collected black spruce boughs up to 30 cm in length and up to 0.5 cm in diameter at the closest tree to the sampling point in each of the NW, NE, SE, SW directions (trees <15 m away from the sampling point). We strived to collect 25% of our harvest at each of the trees identified around the sampling point. Spruce boughs were collected only from mature trees (>2 m), as they are more palatable to hares and are less likely to cause significant weight loss during feeding trials than boughs from juvenile trees (Rodgers and Sinclair 1997). We only collected boughs with healthy needles, and from branches < 1.5 m from ground height, to harvest what would be accessible by a snowshoe hare from the summer to winter months, when snow allows access to higher branches. Although the individual trees that were sampled to inform the stoichiometric distribution map at this grid were juveniles, mature trees have shown similar stoichiometric trends as juveniles (Balluffi-Fry et al. 2022). We assumed that the % N in the spruce at the time we harvested was still representative of the %N during sampling (Heckford et al. 2021), based on recent evidence of a lack of interannual differences in %N (Richmond et al. 2020). We stored harvested spruce outdoors to keep it fresh, and boughs were used for feeding trials within six days of collection. Before preparing black spruce for feeding trials, we thoroughly mixed high-quality spruce boughs to ensure a homogeneous distribution of the boughs sampled at each of the four trees of the sampling point and did the same for the low-quality spruce. Boughs were reduced to 10 cm twigs to fit in feeding baskets.
